# Peripheral Blood Immune Cells from Individuals with Parkinson’s Disease or Inflammatory Bowel Disease Share Deficits in Iron Storage and Transport that are Modulated by Non-Steroidal Anti-Inflammatory Drugs

**DOI:** 10.1101/2024.08.19.608634

**Authors:** MacKenzie L. Bolen, Beatriz Nuñes Gomes, Blake Gill, Kelly B. Menees, Hannah Staley, Janna Jernigan, Malú Gámez Tansey

**Affiliations:** Center for Translational Research in Neurodegenerative Disease, College of Medicine, University of Florida, Gainesville, FL, USA; Department of Neuroscience, College of Medicine, University of Florida, Gainesville, FL, USA; McKnight Brain Institute, University of Florida, Gainesville, FL, USA; Aligning Science Across Parkinson’s (ASAP) Collaborative Research Network, Chevy Chase, MD, USA; Department of Surgery, Northwestern University; Norman Fixel Institute for Neurological Diseases, University of Florida, Gainesville, FL, USA

## Abstract

Parkinson’s Disease (PD) is a multisystem disorder in which dysregulated neuroimmune crosstalk and inflammatory relay via the gut-blood-brain axis have been implicated in PD pathogenesis. Although alterations in circulating inflammatory cytokines and reactive oxygen species (ROS) have been associated with PD, no biomarkers have been identified that predict clinical progression or disease outcome. Gastrointestinal (GI) dysfunction, which involves perturbation of the underlying immune system, is an early and often-overlooked symptom that affects up to 80% of individuals living with PD. Interestingly, 50-70% of individuals with inflammatory bowel disease (IBD), a GI condition that has been epidemiologically linked to PD, display chronic illness-induced anemia — which drives toxic accumulation of iron in the gut. Ferroptotic (or iron loaded) cells have small and dysmorphic mitochondria—suggesting that mitochondrial dysfunction is a consequence of iron accumulation. In pro-inflammatory environments, iron accumulates in immune cells, suggesting a possible connection and/or synergy between iron dysregulation and immune cell dysfunction. Peripheral blood mononuclear cells (PBMCs) recapitulate certain PD-associated neuropathological and inflammatory signatures and can act as communicating messengers in the gut-brain axis. Additionally, this communication can be modulated by several environmental factors; specifically, our data further support existing literature demonstrating a role for non-steroidal anti-inflammatory drugs (NSAIDs) in modulating immune transcriptional states in inflamed individuals. A mechanism linking chronic gut inflammation to iron dysregulation and mitochondrial function within peripheral immune cells has yet to be identified in conferring risk for PD. To that end, we isolated PBMCs and simultaneously evaluated their directed transcriptome and bioenergetic status, to investigate if iron dysregulation and mitochondrial sensitization are linked in individuals living with PD or IBD because of chronic underlying remittent immune activation. We have identified shared features of peripheral inflammation and immunometabolism in individuals living with IBD or PD that may contribute to the epidemiological association reported between IBD and risk for PD.

## INTRODUCTION

Parkinson’s Disease (PD) is a spectrum disorder characterized by an array of both early non-motor (sleep disturbances, anosmia, depression, GI dysfunction, etc.) and motor (bradykinesia, rigidity, postural instability, dystonia, etc.) symptoms. Following the initiation of motor symptomatology, it is estimated that between 50-70% of the dopaminergic (DA) neurons within the substantia nigra pars compacta (SNpc) have been lost^1^. Identifying early and easily detectable biomarkers before irreversible DA cell death occurs is critical for advancing timely therapeutic strategies.

Recent epidemiologic studies have reported increased risk of PD in individuals diagnosed with inflammatory bowel disease (IBD). IBD is characterized by a feed-forward inflammatory response in the gut microenvironment and gut dysbiosis, catalyzing a remittent inflammatory immune response^2^. As IBD has been epidemiologically associated with increased risk for PD, we hypothesize that chronic gut inflammation contributes to ‘body-first’ PD which may initiate in the gut decades prior to the development of classical PD motor symptoms^3–4^. However, currently there is no clear molecular mechanism linking PD risk in individuals living with IBD; and the role of the overactive immune system as the feed-forward inflammatory highway connecting the two diseases remains to be fully understood. We posit that the peripheral immune system can act as a communicating messenger between the inflamed gut and brain and hypothesize the existence of an interface where in the context of GI dysfunction, circulating blood immune cells communicate inflammatory signals to the brain and contribute to increased risk for neurodegenerative disease.

Tissue inflammation, blood loss, and an increase in inflammatory cytokines leads to 79-100% of individuals living with IBD developing anemia at least once^5–6^ making anemia the most frequent metabolic complication of IBD^6^. Inflammation-induced anemia or anemia of chronic disease (ACD) drives a cytotoxic increase of intracellular iron^7^. Iron is a tightly regulated micronutrient that is fundamental to life sustaining biochemical processes such as energy transport, immune cell and mitochondrial function.^8–9^ It is known that infiltrating immune cells are critical in neuroinflammation. Interestingly, these same infiltrating immune cells have been reported to have excess ferritin (the primary measure of iron storage),^10–12^ indicating a synergy between toxic iron accumulation and inappropriate immune cell activation. Iron trafficking is regulated by cytokine interaction^13–15^; therefore, since IBD induces explosive phases of cytokine storms, IBD can lead to critical dysregulation of iron import and export receptors.^16^ Furthermore, the gut microbiota can directly dysregulate the storage and trafficking of iron.^17^ Iron loading as a result of ACD impairs complex I and II in the electron transport chain (ETC) within immune cells which induces excess ROS production,^18^ immune cell activation and proliferation.^13^ Excess ROS produced by increased ferric iron content in the brain induces DA neuron degeneration in mice and rats^19^ and additional work has linked the accumulation of ferric iron to pathologic aggregation of several neurodegenerative-associated proteins, including α-synuclein, tau, amyloid beta (Aβ), and TDP-43.^19^ Several papers have confirmed the role of iron in immune activation and resulting neurodegeneration13.^13, 20–21^ Additionally, several reports have identified individuals with hereditary peripheral iron overload, hemochromatosis, have an increased risk for idiopathic PD,^22–25^ indicating a direct link between iron metabolism and PD etiopathogenesis.

The gut houses up to 80% of the body’s immune cells and is considered the largest immune-pathogen interface in the body.^26^ In the context of dysbiosis, pathogenic bacteria can permeate the gut barrier and catalyze a systemic immunogenic response.^27–28^ Circulating peripheral blood mononuclear cells (PBMCs) are abundant and an easily accessible biosample that can be leveraged to assess immune-related phenotypes of disease risk and progression. PBMCs provide a window into the brain as they recapitulate gene expression seen in postmortem CSF and parallel transcriptional editing seen in neurons from individuals living with PD.^29–30^ Communication between circulating immune cells, like PBMCs, and gut microbiota and their metabolites protect the host from pathogen invasion.^31^ We hypothesize that this disease-related immune signature begins with gut dysbiosis (Fig 1). However, the simultaneous assessment of transcriptional and bioenergetic profiles of PBMCs from individuals with PD or IBD has yet to be published. Therefore, we isolated PBMCs from PD, IBD and neurologically healthy control (NHC) whole blood samples to further investigate transcriptional profiling and bioenergetic functional analyses. Due to the critical role systemic inflammation plays in the regulation of both iron and health of mitochondrial function, we also investigated the impact that anti-inflammatory medication may play on circulating immune cell fitness. Non-steroidal anti-inflammatory drugs (NSAIDs) have been associated with a decreased risk in development of several neurodegenerative diseases.^32–33^ In the context of rodent models, NSAIDs protect DA neurons in vitro against MPTP-induced neurotoxicity.^34–35^ Therefore, we investigated the extent to which self-reported NSAID regimens modulated the inflammatory blood cell immune phenotypes in individuals with IBD or PD compared to NHC.

**Figure 1.**
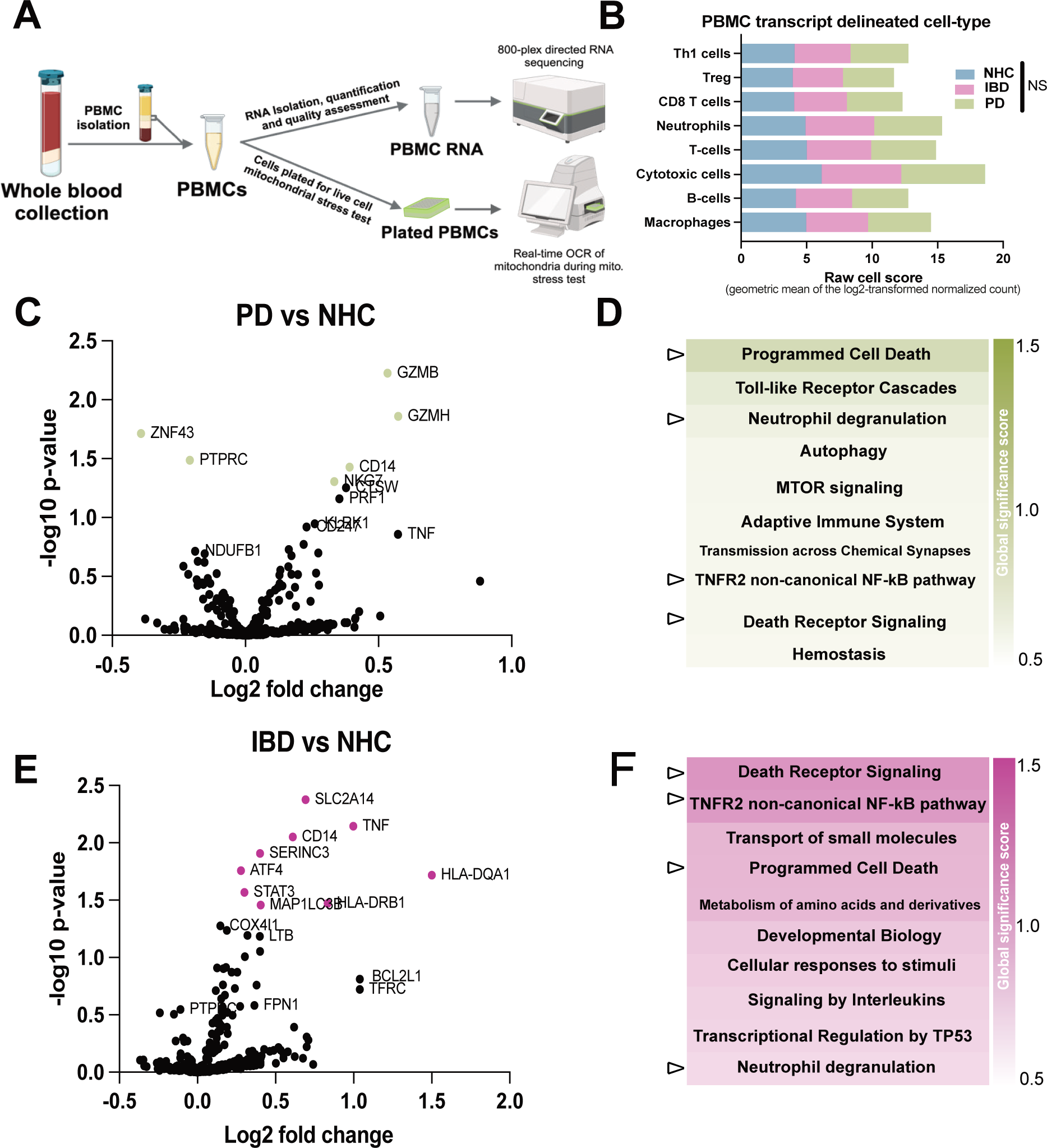
PD and IBD PBMCs display shared and disease-specific transcriptional profiles compared to NHC. **A)** Collection of whole blood and isolation of PBMCs for downstream directed transcriptomic and real-time mitochondria stress test. **B)** Cell phenotyping represented and quantified by the raw cell count score, generated via the geometric mean of the Log2 normalized count. **C, E)** PD and IBD PBMC mRNA transcript counts are compared against NHC and each other. Volcano plots indicate transcripts that are significantly enriched or depleted (green, purple or blue significance indicated as *p* <0.05). **D and F)** Top 10 pathways reflecting the impact of differential expression within each gene set pathway when comparing PD to IBD PBMC, quantified via the global significance score calculated via the square root of the mean squared t-statistic for the genes in a gene set. A larger global significance score indicates more DEGs within that pathway. Shared enriched pathways between PD and IBD PMBCs (as compared to NHC) are indicated with ⊳.

## METHODS

### Human Subjects

This study was reviewed and approved by the University of Florida Institutional Review Board (IRB201902459). Informed consent was required for participation. In a private area, the research coordinator thoroughly explained the study and the individual’s involvement. If the individual elected to participate in the study, they signed the consent, and all participants signed an informed consent, approved by UF IRB. Individuals living with PD were recruited from the University of Florida Neuromedicine Clinic at the Fixel Institute for Neurological Diseases. Individuals living with IBD who were already preparing for routine colonoscopy were recruited from the UF Gastrointestinal (GI) Clinic. Neurologically Healthy control individuals were all PD-spousal controls except for one subject that was an IBD spousal control. Participation within this cross-sectional study included inclusion criteria of reported age between 40-80 years, diagnosis of PD (based on MDS score), IBD, or no diagnosis in the case of the neurologically healthy control cohort. Individuals were excluded from the study if those diagnosed with PD reported active infections or active autoimmune or chronic inflammatory conditions that required treatment with immunosuppressants. Additional exclusion criteria applicable included pregnancy, history of a blood transfusion within 4-weeks of recruitment, body weight less than 110 lbs., subjects who were perceived by the investigator to be unable to comply with study procedures, and treatment with immunosuppressants or antibiotics within one month of recruitment. Notably, all IBD subjects recruited for this study were declared to be in remission, as confirmed by colonic biopsies that were collected by the gastroenterologists as part of routine colonoscopies.

Blood was collected from 17 individuals with PD, 15 individuals with IBD, and 15 neurologically healthy controls. A total of 24 individuals identified as biologically female and 23 identified as biologically male. The neurologically healthy controls were spousal controls of the individuals with PD. A questionnaire to collect information about age, sex, race, lifestyle habits such as smoking, over-the-counter or prescription NSAID use, and probiotic use was completed by all participants. Health history was collected as well, including assessment of disease duration, MoCA and Schwab scores, past or current endocrine dysfunction, and cancer history. Due to minimal representation of non-white ethnic groups within our study and the fact that ethnicity influences immune system responses, we limited our analyses to samples from individuals who self-identified as “white”. Future studies will focus on recruitment of a more heterogenous population.

### Peripheral blood mononuclear cell Isolation

PBMCs were harvested as previously published.^36^

### Genomic DNA Isolation and Genotyping

Because mutations in the gene encoding for the Leucine-rich repeat kinase 2 (*LRRK2*) are the most common genetic cause of autosomal dominant familial and sporadic PD and IBD, we performed genotyping analysis to exclude any individuals with a *LRRK2* mutation or variant associated with the latter. Genomic DNA (gDNA) isolation was performed on all PBMC samples using the DNeasy Blood and Tissue Kit (QIAgen, 69504) according to manufacturer’s instructions. The final flowthrough was quantified using the DeNovix DS-11 Spectrophotometer and the samples were diluted with DNAse-free water to ensure the presence of 50ng of DNA in each final well. The genotyping working mixture was prepared by combining 0.5uL of 40X custom SNP primer (ID: rs34637584) for LRRK2:G2019S, 0.5uL of 1X TE buffer, and 11uL of TaqmanTM Fast Advanced Master Mix (AppliedBiosystemsTM, 4444556). Subsequently, 11uL of this working mixture was added to 9uL of the diluted gDNA. In triplicate, 5ul of this final mixture was dispensed into a 384-well plate. Following centrifugation, the plate was loaded into the QuantStudio5 qPCR machine, and the PCR cycle was initiated. This cycle included a pre-read at 60°C for 30 seconds, an initial hold at 95°C for 10 minutes, followed by 40 PCR cycles at 95°C for 15 seconds and 60°C for 1 minute each. A post-read was performed at 60°C for 30 seconds and placed on hold at 4°C. Two participants self-reported as LRRK2:G2019S mutation positive and were used as positive controls for the genotyping analysis.

### XF96 Seahorse flux Analyzer

The XF96 Seahorse flux Analyzer (Agilent, Santa Clara, CA, USA) was employed to monitor mitochondrial bioenergetic capacity in real-time in PBMCs from the PD, IBD, and NHC cohorts. Parameters such as oxygen consumption rate (OCR) and extracellular acidification rate (ECAR) were assessed following sequential injections of oligomycin, carbonyl cyanide-4 (trifluoromethoxy) phenylhydrazone (FCCP), and a combination of antimycin A and rotenone. PBMCs were stained with Hoechst 33342 and imaged, with cell numbers in each well used to normalize all OCR parameters.

### RNA Isolation

RNA was extracted from freshly-thawed PBMCs using the Qiagen RNeasy kit (Qiagen, Valencia, CA) (CAT# NC9307831) according to manufacturer’s instructions on the same day that XF96 Seahorse cell stress test was completed and from the same sample vile. The final flowthrough was quantified using the DeNovix DS-11 Spectrophotometer and stored at −80°C until analysis.

### NanoString nCounter Transcriptional Analysis

The NanoString nCounter® Analysis System (NanoString Technologies, Seattle, USA) was employed to conduct the Human Cell Metabolic Pathways nCounter® assay (CAT# XT-115000367). This assay utilized a panel consisting of 760 target genes, accompanied by an additional set of 10 internal reference genes. These internal reference genes, namely *CYTB, DMT1, DUOXA2, IREB2, LAMP2, FERT, FPTN1, HAMP, MCP1,* and *RAB8*, functioned as custom spike-in panels to ensure quality control and enable normalization of the data.

Upon retrieval from −80°C storage, samples were thawed on wet ice, and RNA concentration and quality were assessed using the Qubit and Quant-it Assay (CAT# Q33263) and Agilent 2100 Bioanalyzer (CAT#G2939A). To standardize the RNA concentration across samples, each sample was diluted with an appropriate volume of nuclease-free water to achieve a final concentration of 100 ng of RNA in 5μL of water. In sequence, the hybridization step was performed following the manufacturer’s instructions. Post hybridization, the samples were pipetted into the nCounter chip via the nCounter MAX/FLEX robot where non-adhered capture and reporter probes were removed and hybridized mRNAs were immobilized for imaging.

### nCounter Data Analysis

Raw count RCC files were normalized on the nSolver analysis software v4.0 (NanoString Technologies) according to the manufacturer’s instructions. Csv files were then exported into Graph Pad PRISM, where built-in statistical analysis was completed per manufacturer’s instructions. Normalization of all gene expressions was performed using NanoString nSolver v4.0 default settings, where raw counts are normalized to the geometric mean of positive/negative controls and housekeeping probes within the panel. Changes in expression ratios were produced by dividing the mean values in the treatment group by the mean values of the reference group. *P* values were considered significant if *p* < 0.05 and the fold change was greater than [1.5]. Adjusted *p-*values were produced from the Benjamini-Hochberg method of false discovery rate (FDR) corrected *p*-values. All raw counts were normalized to all 10 reference genes. Probes were not selected for analysis if greater than half of the samples did not meet the sample-specific threshold, as a result, said pruned probes produced a log2 Fold Change of 0 and an adjusted *p* value of 1. Soft independent modeling by class analogy (SIMCA) (v18.0., Item No.: UT-SS-1232, Umetrics, Umea, Sweden) was leveraged to produce biplots visualizing multivariate analysis of principle components and variance within these data (Fig 2A).

**Figure 2.**
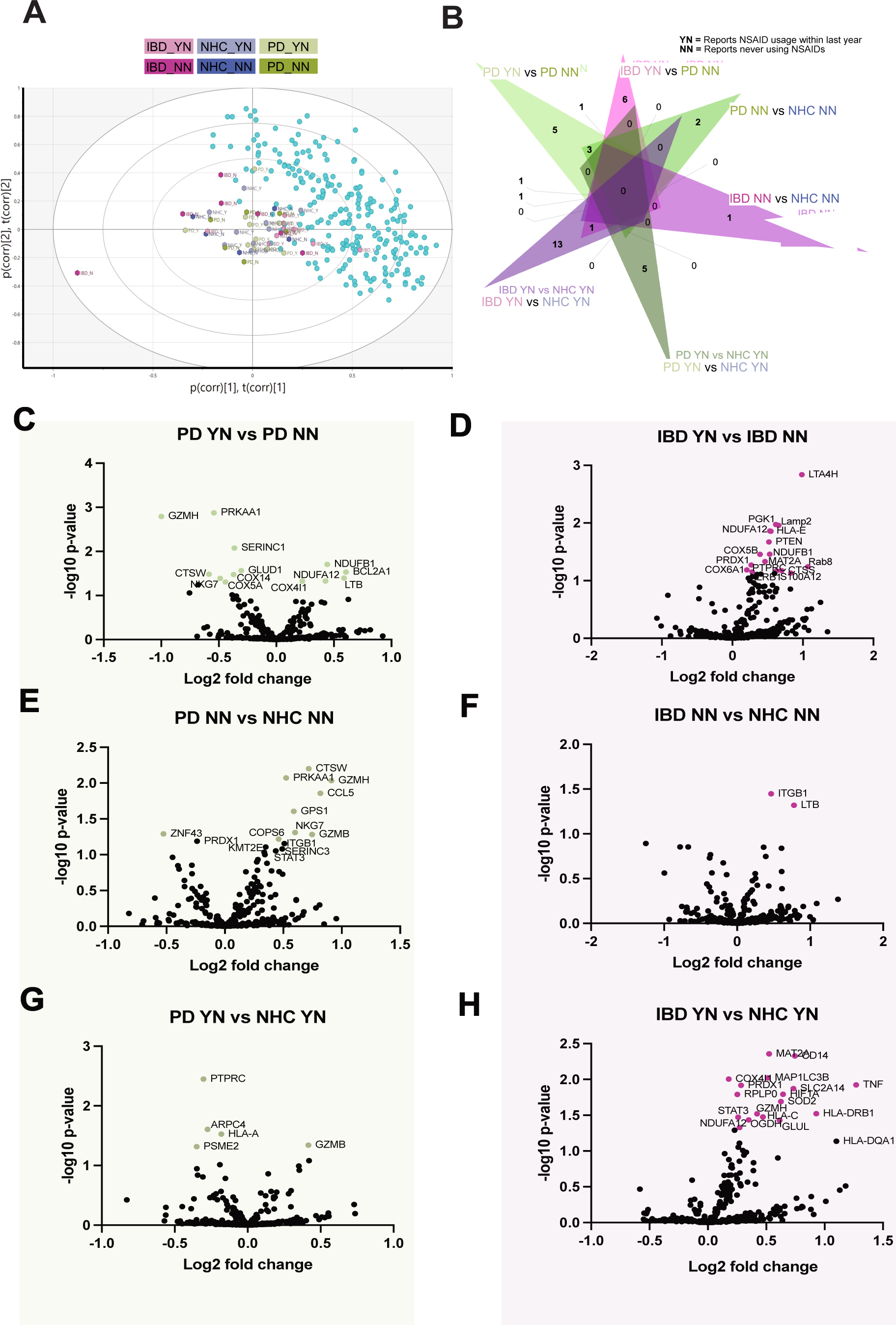
Self-reported NSAID use modulates disease-dependent transcriptional profiles of PBMCs. **A)** Principal components analysis was completed via Soft independent modeling by class analogy (SIMCA) (v18.0, Umetrics, Umeå, Sweden) to produce biplots visualizing multivariate analysis of transcripts by disease and reported NSAID use **B)** Using jVenn, a 6 group Venn diagram was generated comparing the significant DEGs by disease and NSAID usage. **C-H)** PD and IBD PBMC mRNA transcript counts are compared against NHC, basedon reported NSAID use or lack thereof. Volcano plots indicate transcripts that are significantly enriched or depleted (green = comparison with PD, purple = comparison with IBD), significance indicated as *p* <0.05 with either green or purple colored dot.

## RESULTS

Herein, we investigated the extent to which 1) IBD and PD PBMC transcriptional profiles share common features as compared to NHC (Fig 1), and the potential modulation of such by anti-inflammatory medication use (Fig 2); 2) IBD and PD PBMC mitochondria have increased oxygen consumption (Fig 4); and 3) the relationship between iron-related transcript regulation and mitochondrial function in PBMCs (Fig 6).

### PD and IBD PBMCs display shared and disease-specific transcriptional profiles compared to NHC

To investigate the directed metabolism-associated transcriptome from 17 PD, 15 IBD and 15 NHC age- and sex-matched individuals (Table 1) we utilized the NanoString nCounter™ platform. After PBMCs were isolated from whole blood (Fig 1A), individual immune cell-types were identified via gene enrichment (B Cells = *BLK, CD19, FAM30A, FCRL2, MS4A1, PNOC, SPIB, TCL1A, TNFRSF17*; CD8 T Cells = *CD8A, CD8B*; Cytotoxic cells = *CTSW, GNLY, GZMA, GZMB, GZMH, KLRB1, KLRD1, KLRK1, NKG7, PRF1*; Dendritic cells = *CCL13, CD209, HSD11B*; Macrophages = *CD163, CD68, CD84, MSFA4A*; Neutrophils = *CEACAM3, CSF3R, FCAR, FPR1, S100A12, SIGLEC5*; T cells = *CD3D, CD3E, CD3G, CD6, SH2D1A, TRAT1*; Th1 cells = *TBX21*; T regulatory cells = *FOXP3*). There was no significant difference between gene-identified cell classification abundance across diseases (Fig 1B).

**Table 1.**
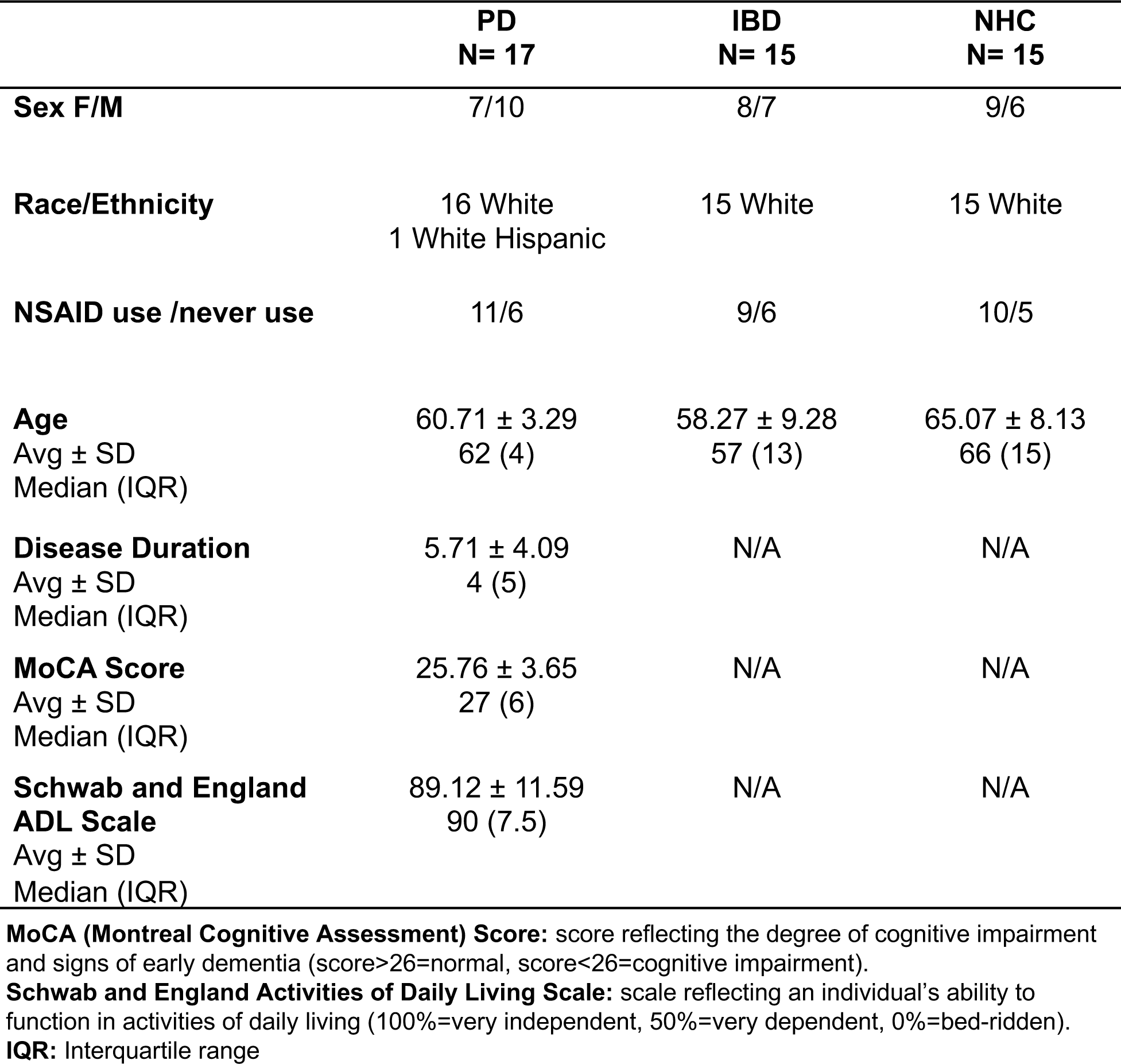
Participant demographics and cognitive status.

Overall, PD PBMCs displayed significant enrichment of transcripts associated with immune activation (i.e. *GZMB/H, CD14*) (Fig 1C). These disease-specific immunoregulatory gene-set outcomes were also characterized by significant changes in pathways associated with the adaptive immune system: TNF and NFκB signaling (Fig 1D). IBD PBMCs displayed a substantial enrichment of inflammatory genes associated with antigen presentation (i.e., *TNF* and *HLA*), compared to NHC PBMCs (Fig 1E). IBD PBMCs also displayed enrichment in inflammatory pathways associated with interleukin signaling and cellular responses to other stimuli (Fig 1F). Of interest were the shared enriched pathways between PD and IBD PBMCs (as compared to NHC), which included death receptor signaling, TNFR2 non-canonical NFκB pathway, programmed cell death, and neutrophil degranulation (Fig 1 D and F).

### Self-reported NSAID use modulates disease-dependent transcriptional profiles of PBMCs

PD and IBD are diseases characterized by chronic underlying inflammation. Due to the extensive literature discussing the effects that anti-inflammatory medications can have on immune cell transcriptional status^37–40^ we stratified PD, IBD and NHC individuals by reported non-steroidal anti-inflammatory drug (NSAID) use. Samples were separated by those having reported never having used NSAIDs (NN) or having reported recent (within the last two weeks of blood collection) and frequent (used 1 or more times per week) use of NSAIDs (YN). Principal components analysis was completed via Soft independent modeling by class analogy (SIMCA) (v18.0, Umetrics, Umeå, Sweden) to produce biplots visualizing multivariate analysis (Fig 2A). We identified significantly different gene expression profiles based on NSAID use and disease status (Fig 2). Interestingly, some DEGs were shared between comparisons, such as a significant increase in *NDUFA12* in both PD YN and IBD YN as compared to both PD NN (Fig 2B, C) and IBD NN (Fig 2B,D). However, when compared to NHC, *NDUFA12* is only enriched in IBD YN vs NHC YN (Fig 2A,F) with no significant difference in *NDUF12A* enrichment in any PD group vs NHC (Fig 2E,G) or IBD NN vs NHC NN (Fig 2F) in PBMC transcript counts. Additionally, *PRKAA1, GZMH* and *CTSW* were all significantly depleted in PD YN when compared to PD NN (Fig 2C). All 3 of these genes were also significantly enriched in PD NN as compared to NHC NN (Fig 2E), indicating an NSAID effect on PBMC transcriptional signatures. Both PD NN and PD YN displayed enrichment of *GZMB* as compared to NHC NN (Fig 2E) and NHC YN (Fig 2G). These data suggest that NSAID uses can modify the transcriptional profiles of PBMCs regardless of disease status.

### NSAID use abolishes differences between PD and IBD inflammation-associated immune profiles

We next compared PD and IBD transcript counts directly (Fig 3A). Transcripts associated with mitochondrial subunit function and regulation were depleted in PD, which is additionally indicated by significant enrichment of global significance scores indicating implicated gene sets in metabolism and respiratory electron transport as compared to IBD (Fig 3B). When comparing PD and IBD PBMCs from individuals who self-report NSAID use, PD PBMCs displayed significant depletion in transcripts associated with antigen presentation, like *HLA-DQA1* and *PTPRC*, which was not present in the comparison of PD vs IBD who do not report chronic NSAID use (Fig 3C). In line with previous work^41^, individuals with PD who report never having used chronic NSAIDs have a significant enrichment of *GZMB* and *PRF1*, both of which are associated with cytotoxic T-cell activation, as compared to individuals with IBD who report not ever having used NSAIDs (Fig 3D).

**Figure 3.**
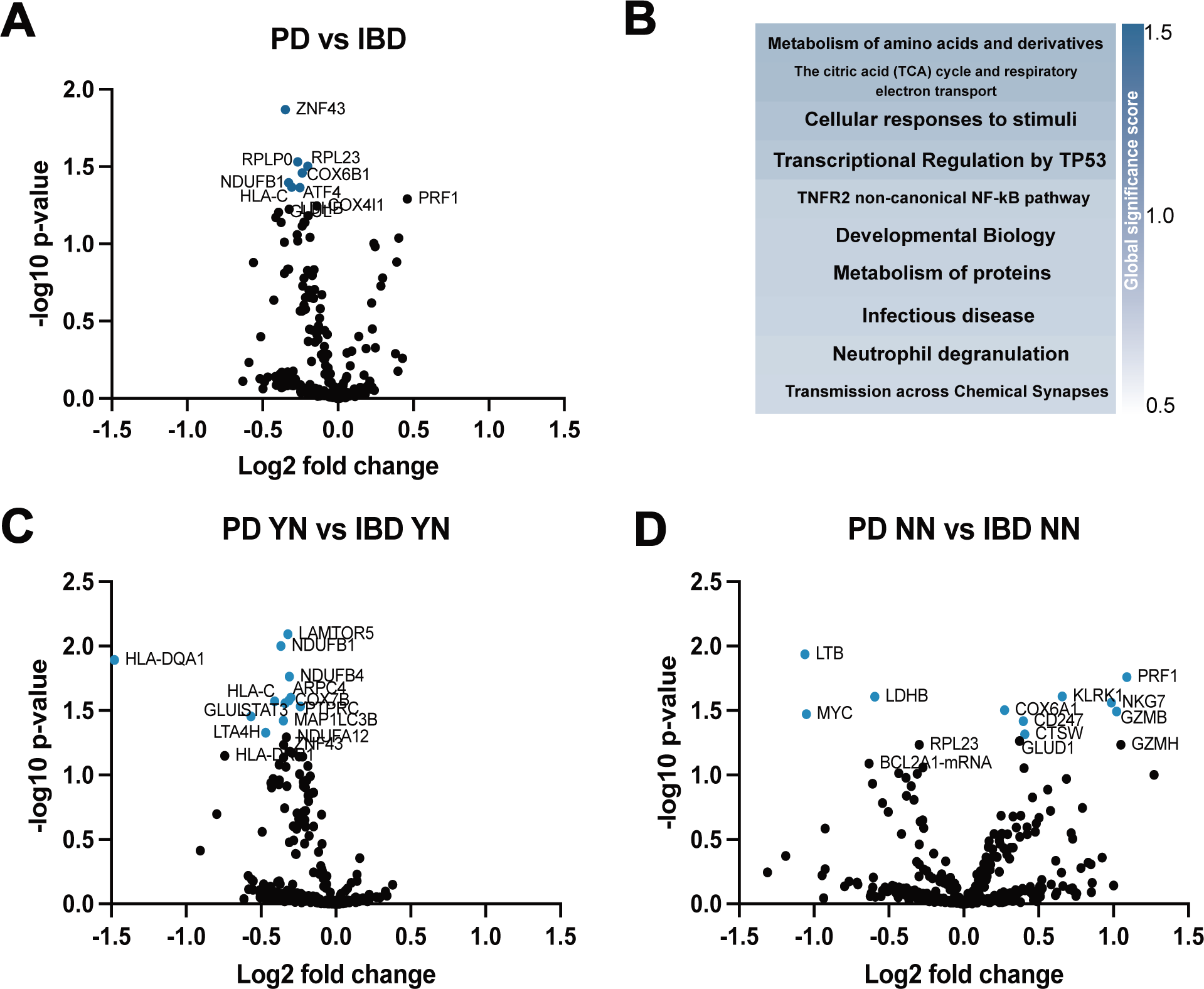
NSAID use abolishes differences between PD and IBD inflammation-associated immune profiles. **A)** PD and IBD PBMC mRNA transcript counts are compared against each other **B)** Top 10 pathways reflecting the impact of differential expression within each gene set pathway when comparing PD to IBD PBMCs, quantified via the global significance score calculated via the square root of the mean squared t-statistic for the genes in a gene set. A larger global significance score indicates more DEGs within that pathway. C) PD and IBD PBMC mRNA transcript counts are compared to each other, based on reported NSAID use or lack thereof. Volcano plots indicate transcripts that are significantly enriched or depleted, significance indicated as *p* <0.05.

### Real-time measurement of mitochondrial oxygen consumption from PD and IBD PBMCs indicates no significant differences in respirometric function

To investigate mitochondrial function in PBMCs in real time, we leveraged the mitochondrial stress test on the Seahorse XF analyzer, which utilizes various mitochondrial complex inhibitors to measure oxygen consumption. When measuring basal, maximal, spare and ATP production based on oxygen consumption of PBMCs, there was no significant difference between diseases or as compared to NHC PBMCs (Fig 4 A,C). However, there were significant differences in the distribution of variance between disease and compared to NHC (Supplemental Fig 1A-D). Although not statistically significant, when separating groups by NSAID use, there was an increase in oxygen consumption by PD and IBD PBMCs from individuals who reported NSAID use as compared to PD and IBD PBMCs from individuals who reported no NSAID use and as compared to NHC (Fig 4 B,D) (Supplemental Fig 1 E-H).

**Figure 4.**
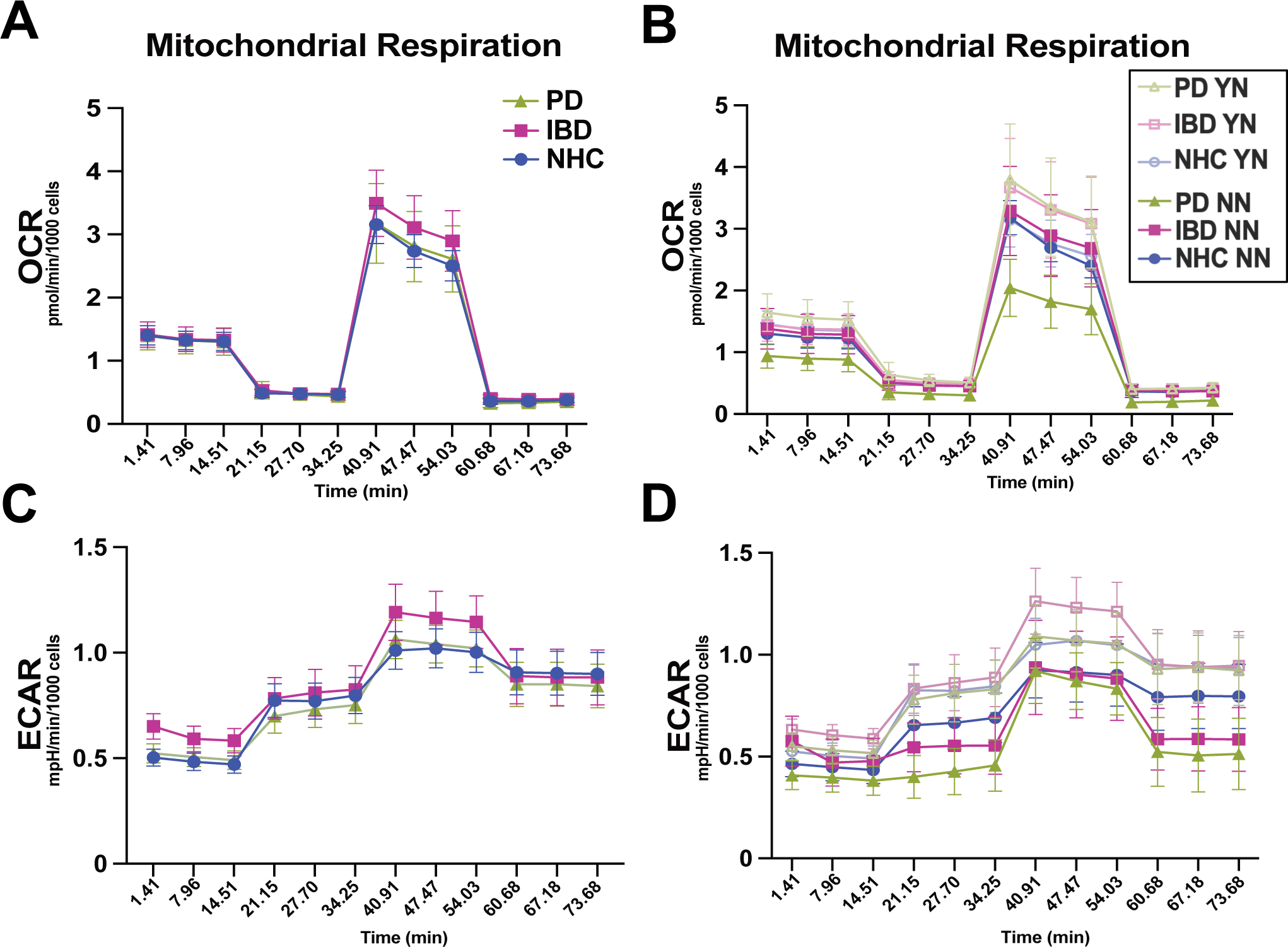
Real-time measurement of mitochondrial oxygen consumption from PD and IBD PBMCs indicates no significant differences in respirometric function. **A-D)** Average trace of oxygen consumption rate from adhered PBMCs over time, all values are normalized to cell count and variance is indicated via ± S.E.M (YN = NSAID user, NN = does not report ever using NSAIDs).

### Enrichment of iron-related transcripts associated with peripheral inflammation have a significant disease-dependent positive correlation with increase in oxygen consumption

An excess of the iron storage protein ferritin and dysregulation of the iron importer and exporters *FPTN-1* and *DMT1* respectively, have been directly linked to mitochondria dysfunction.^8–9^ To investigate the relationship between enrichment in iron metabolism transcripts and mitochondrial oxygen consumption, we preformed linear regressions and found several positive significant relationships between enrichment of iron-associated transcript based on phase of mitochondrial oxygen consumption and disease (Fig 5). We observed that, during basal respiration (baseline consumption of oxygen for mitochondria), both PD and IBD PBMCs that consume more oxygen at baseline also have a higher ferritin content (Fig 5). We also preformed linear regressions of oxygen consumption relative to iron-associated transcripts on PD and IBD PBMCs relative to NHC, grouped by NSAID use (Supplemental Fig 2) and found NSAID dependent correlations between iron metabolism and mitochondrial oxygen consumption.

**Figure 5.** Enrichment of iron-related transcripts associated with peripheral inflammation have a significant disease-dependent positive correlation with increase in oxygen consumption. A star indicates a significant positive relationship between increase in transcript count and increase in the oxygen consumed during the specific mitochondrial phase, the R^2^ correlation is graphed to 1 and indicated by an increase is blue opacity, significance is indicated with * = *p* < 0.05, ** *p* <0.001, *** = *p* <0.0001.

## DISCUSSION

The aim of these experiments was to investigate the impact of disease-associated inflammation in IBD and PD on PBMC immune activation and mitochondrial respiration. These data revealed 3 novel findings: 1) PD and IBD PBMCs share several inflammatory signatures compared to NHC and the specific transcripts enriched or depleted are disease-dependent (Fig 1); 2) The effect of chronic NSAID use on PBMC transcriptional profiles is robust and disease independent (Fig 2); 3) PD and IBD PBMCs both display enrichment of iron-related transcripts with an increase in specific OCR phases (Fig 5).

In support of our hypothesis, we observed that the increased frequency in iron-transcript dysregulation in IBD and PD PBMCs but not in NHC PBMCs directly correlates with a change in mitochondrial function. Excess ferric iron is housed in ferritin, which is the dominant ferric iron storage protein in the central nervous system and a surrogate for total-iron content.^42^ In individuals with PD, there is an increase in ferritin load in the cerebrospinal fluid (CSF) as compared to age-matched controls.^42^ A separate study also reported that ferritin concentrations increased in PD CSF with progression of motor symptom severity.^43^ Increases in iron deposition have been reported in mitochondrial dysfunction and can catalyze an inflammatory phenotype in immune cells^10^– which our findings suggest may be abolished with chronic use of NSAIDs.

Our study is the first to directly compare the metabolism-associated transcriptional profiles of PBMCs from individuals with PD and IBD. Our data suggest that both PD and IBD cohorts are peripherally inflamed, and these profiles of peripheral inflammation are detectable at the transcript level in PBMCs. These data also indicate that although PBMC transcript phenotype is a consequence of disease, the abundance and frequencies of different immune cell-types did not significantly change as a function of disease. By investigating the transcriptional topography of PBMCs from both individuals with IBD and PD we identified that PBMCs reflect a disease-dependent profile that is shifted when exposed to NSAIDs. Our data support the idea that NSAID usage by individuals with PD and IBD aids in suppressing the overactive immune system; however, the effect of these anti-inflammatory drugs on the immune environment appears to be disease dependent (Fig. 3). These data corroborate previous epidemiological studies stating that common use of anti-inflammatories decreases risk for PD in individuals with IBD.^43^ On the basis of our novel findings and other existing data, we propose a model to describe the effects of chronic inflammation associated with IBD or PD on the gut-blood-brain axes. Specifically, where chronic remittent proinflammatory immune activity in the intestine in both individuals living with IBD or PD affects peripheral blood immune cells which act that carry critical messages of inflammatory injury from the periphery to the brain and vice versa.

Peripheral mitochondrial dysfunction has been heavily implicated in PD pathogenesis.^43–44^ In the 1980’s, the famous California case of accidental drug use of 1-methyl-4-phenyl-1,2,3,6-tetrahydropyridine (MPTP), an analog of heroin resulting from sloppy chemistry and a specific DA neuron mitochondrial complex 1 inhibitor, marked a significant breakthrough in understanding PD pathophysiology. Within days of such MPTP-contaminated heroine developed features of parkinsonism and *post-mortem* analysis revealed severe lesioning of DA neurons within the SNpc.^45^ Additionally, rodent models of mild systemic mitochondrial dysfunction provide substantial evidence that peripheral mitochondrial complex damage or respirometric capacity challenge is implicated in the progression of neuroinflammation.^46^ MPTP induces an immunometabolic activation that catalyzes mitochondrial danger-associated molecular pattern (DAMP) release, including reactive oxygen species (ROS) production which results in an increase of cytokine release by neighboring cells and triggers a feed-forward inflammatory response.^47^ There is discordant literature on the extent to which PBMCs from individuals with idiopathic PD have dysregulated mitochondrial respiration. While some reports have identified an increase in maximal and spare respiratory capacity in PD PBMCs^45^, other studies report no difference in any phase of mitochondrial oxygen consumption in PBMCs from individuals with PD as compared to NHC or those with rapid-eye movement (REM) behavior sleep disorder (RBD), a common feature in prodromal PD.^48^ Using linear regression analyses we correlated increase in oxygen consumption separated by mitochondrial respiratory phase with an increase in transcript count associated with iron metabolism. These outcomes suggest an increase in overall cell fitness and energetic capacity that may represent an early-stage compensatory mechanism in response to inflammation-triggered loading of mitochondria in individuals with IBD or PD. We believe that PBMCs reveal system-level peripheral inflammation and immune cell iron regulation, thereby reflecting the well-established ferroptotic cell stress in the brain of individuals living with PD. We were underpowered to parse groups apart by NSAID use within this analysis. However, we identified a unique significant relationship whereby PBMCs from individuals with PD or IBD that displayed enriched ferritin also showed an increase in basal mitochondrial respiration not observed in NHCs. Also, via SIMCA analysis, we identified that ferritin transcript is highly expressed in PBMCs and was one of the most reliably detected transcripts within these data—indicating that ferritin transcripts in PBMCs may be an accessible potential biomarker of immune dysfunction and risk for PD (Supp. Fig 3).

PD risk factors include both environmental and genetic components, with approximately 5-10% of all PD stemming from some form of monogenic familial PD mutation.^49^ The most common genetic cause associated with both autosomal dominant familial and sporadic PD are mutations in the gene encoding Leucine-rich repeat kinase 2 (*LRRK2*), where several pathogenic mutations on this gene are implicated in familial PD including LRRK2 G2019S. Coincidently, specific *LRRK2* mutations are also correlated with increased risk of Crohn’s Disease, a subtype of IBD, thereby implicating LRRK2 as a genetic regulator of inflammatory processes in gut inflammation in both IBD and PD.^51^ For this reason, we genotyped all our samples for the *LRRK2 G2019S* mutation and excluded individuals positive for the mutation from the analysis herein.

Our novel findings support the potential that *in vitro* transcriptional profiling and *in vivo* bioenergetic functional analysis of PBMCs could contribute to elucidation of the mechanisms underlying epidemiological relationships, such as that reported between IBD and risk for PD. Our findings lay the foundation for future mechanism-based rationale to target peripheral immune cell dysfunction to mitigate neuroinflammation and risk for PD in chronic peripheral diseases like IBD.

Although these data provide a unique perspective on the correlational relationship between mitochondrial function and iron transcript regulation, there are several limitations to note. First, this cohort size is small, and these data will require future replication with a larger sample size. Additionally, we collapsed both ulcerative colitis and Crohn’s disease (two types of IBD) cohorts (all individuals with diagnosed IBD were in remission based on pathological analysis of the colonic biopsies) together into a single group, due to only having 3 individuals living with UC, 11 with CD and 1 individual with “both UC and CD”. We detected substantial variation within groups in the seahorse dataset, a larger study may be successful in overcoming the variability in patterns of functional mitochondrial oxygen consumption based on disease and NSAID usage (Supplemental Fig. 1). While we noted hemostasis pathway enrichment in PD PBMCs, this is likely an effect of levodopa usage^52^, yet underscores the importance of taking into account the impact that medication has on functional and transcriptional immunometabolism.

In sum, our studies on transcriptional and functional respirometric profile of peripheral blood immune cells from individuals with IBD, PD and NHC lays the groundwork for additional studies of PBMCs as potentially useful cellular biomarkers of disease risk and progression. Our findings corroborate an overactive immune profile in PBMCs from individuals living with PD^53^ and parse apart possible transcriptional and functional mitochondrial evidence linking IBD as one of many catalysts for development of PD. We identified a correlational relationship between individuals living with PD, whose PBMCs display increased ferritin transcript content with paired increase in mitochondria respirometric capacity in all four functional stages of the ETC but found disparate results in IBD and NHC (Fig 5). Additionally, this investigation further supports the critical need to understand the impact of anti-inflammatories on immunometabolism and how this widely-used drug class could possibly be leveraged as a tool to modulate the inflammatory processes that contribute to pathogenesis of neurodegenerative onset in PD.

## Acknowledgements

We thank members of the Tansey lab for useful discussions. We would also like to thank study participants, without their generous donations none of this work would have been possible.

