## Supplemental Figures for "Peripheral Blood Immune Cells from Individuals with Parkinson’s Disease or Inflammatory Bowel Disease Share Deficits in Iron Storage and Transport that are Modulated by Non-Steroidal Anti-Inflammatory Drugs"

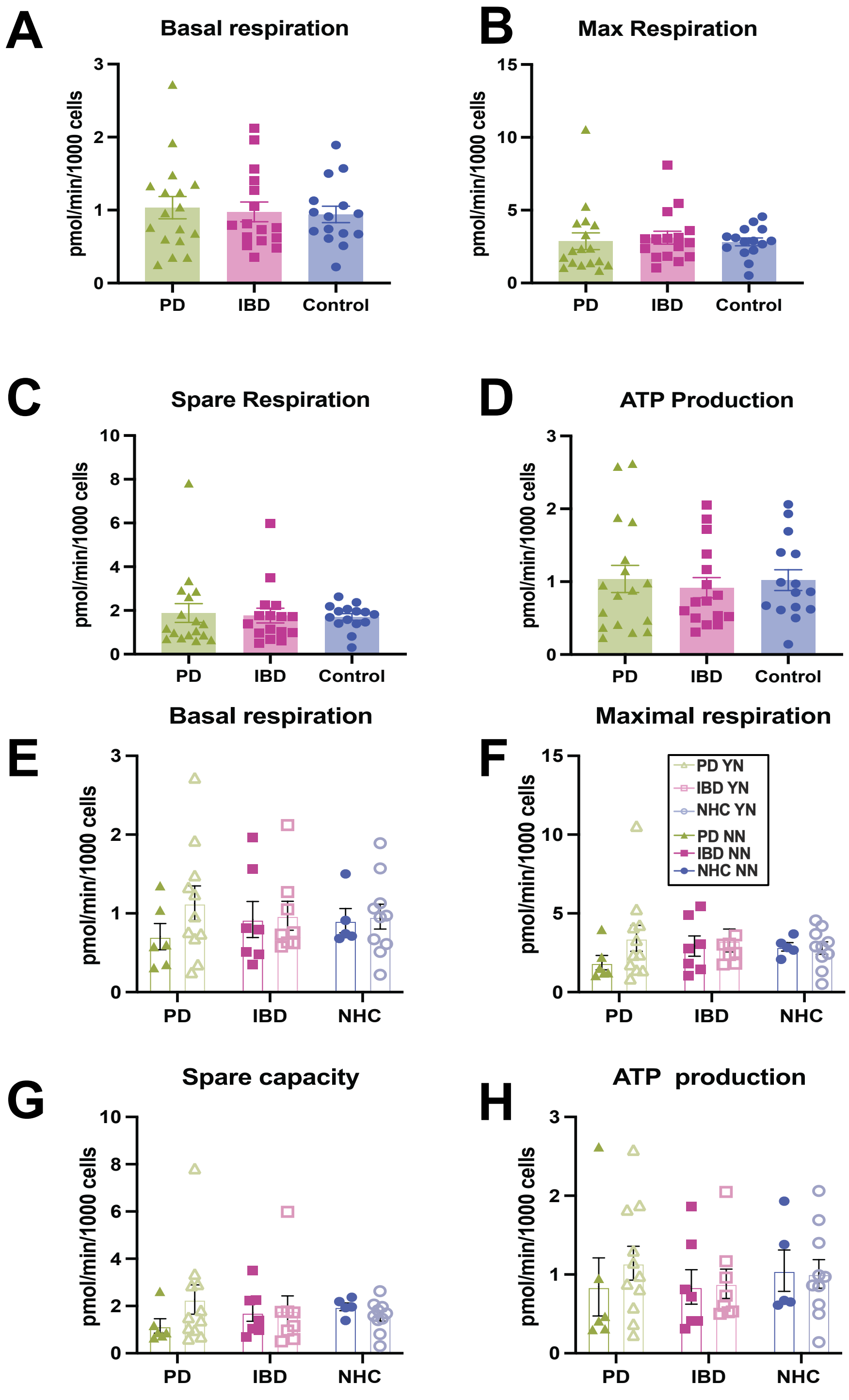


**Supplemental Figure 1. PBMCs from PD, IBD and NHC do not have significantly different oxygen consumption when compared across diseases and to NHC. A-D)** Average trace of oxygen consumption rate from adhered PBMCs over time, all values are normalized to cell count and variance is indicated via ± S.E.M. **E-H)** Average oxygen consumption rate of each mitochondrial respirometric phase parsed apart by NSAID use and disease. No groups were significantly different by two-way ANOVA. YN = NSAID user, NN = does not report ever using NSAIDs. Dark green solid triangle = PBMCs from PD patients who do not report NSAID use. Light green open triangle = PBMCS from PD patients who do report NSAID use. Dark pink solid square = IBD patients who do not report NSAID use. Light pink open square = IBD patients who report NSAID use. Dark blue solid circle = NHC who do not report NSAID use. Light blue open circle = NHC who do report NSAID use.

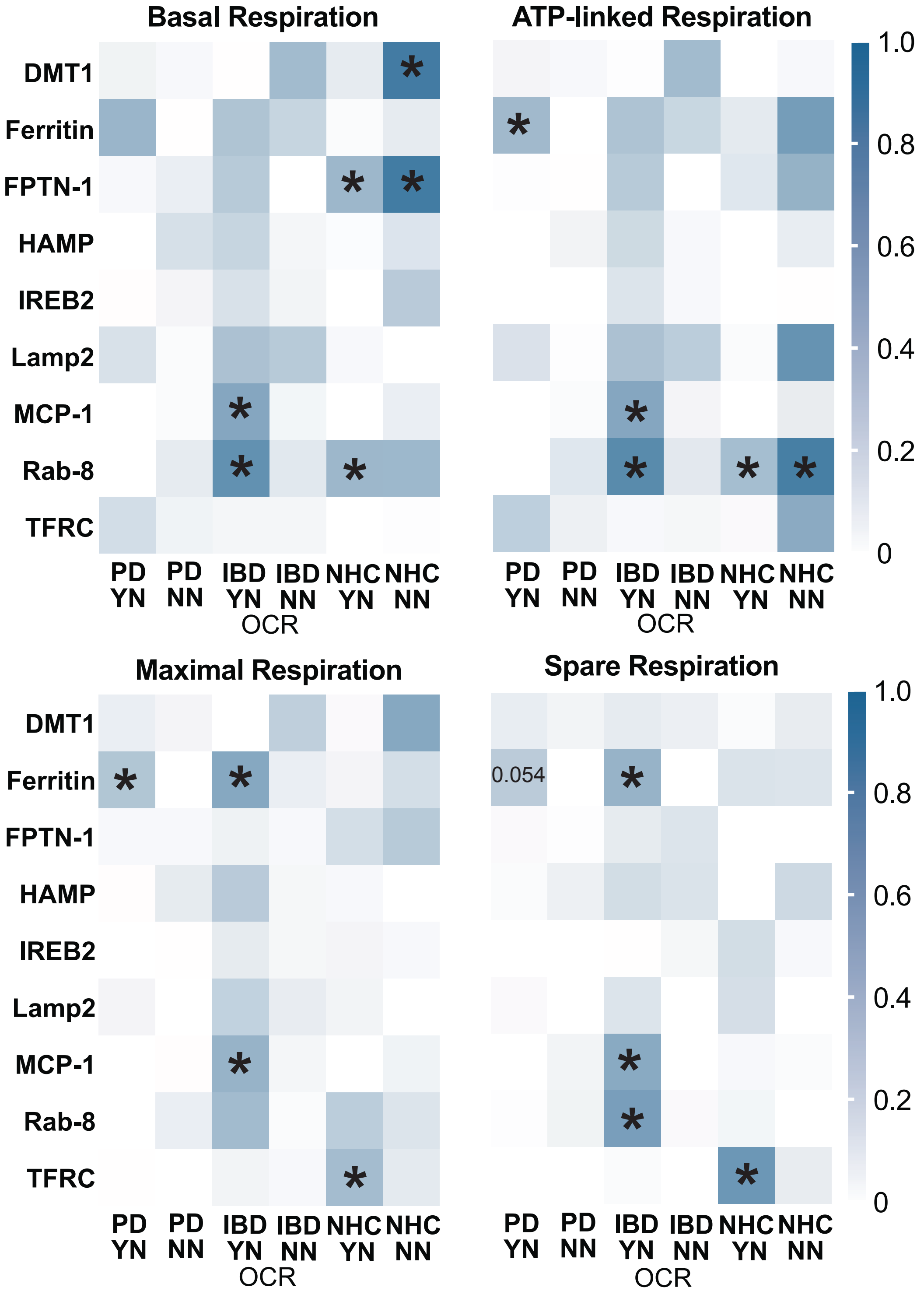


**Supplemental Figure 2. PBMCs indicate a disease and NSAID use dependent co-expression of enrichment in iron-associated genes and increase in OCR across multiple critical parameters of mitochondrial function.** A star indicates a significant positive relationship between increase in transcript count and increase in the oxygen consumed during the specific mitochondrial phase, the $R^2$ correlation is graphed to 1 and indicated by an increase is blue opacity, significance is indicated with * = $p < 0.05$, ** $p < 0.001$, *** = $p < 0.0001$.

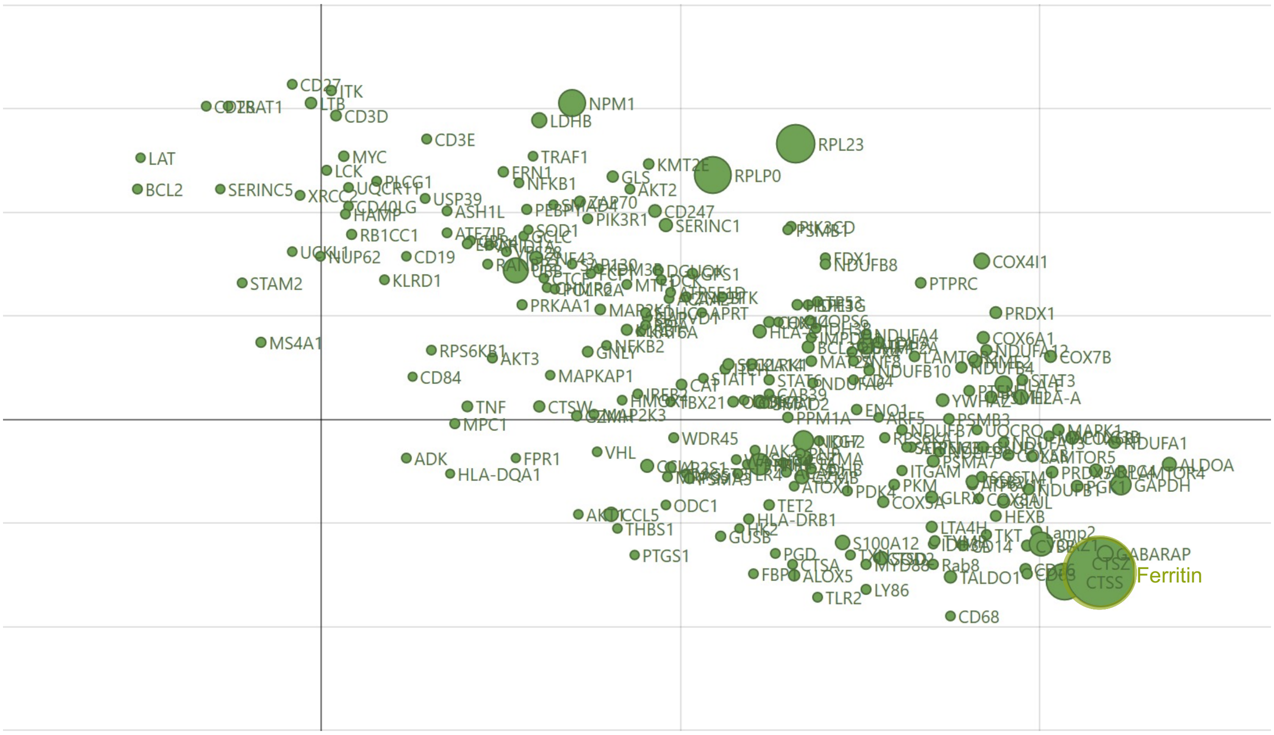


**Supplemental Figure 3. Number of transcript counts within all PBMCs indicate ferritin as the most frequently counted transcript within these data.** Transcripts, which are deemed as loadings in the SIMCA-PCA-X model, are displayed and sorted by vector size where circle size is synonymous to total normalized transcript count. These data were generated using soft independent modeling by class analogy (SIMCA) (v18.0., Item No.: UT-SS-1232, Umetrics, Umeå, Sweden) to produce plots visualizing count analysis of transcripts.
